# AI-based Psychiatric Prediction in Youth: Neuroimaging Provides Minimal Gains Beyond Confounds

**DOI:** 10.64898/2026.05.22.727174

**Authors:** Sam Gijsen, Ammar Ibrahim, Marija Tochadse, Kerstin Ritter

**Affiliations:** Hertie Institute for AI in Brain Health, University of Tübingen, Germany; Tübingen AI Center, University of Tübingen, Germany; Charité – Universitätsmedizin Berlin, Department of Psychiatry and Psychotherapy, Germany; German Center for Mental Health (DZPG), partner site Tübingen, Germany

## Abstract

Recent advances in artificial intelligence (AI) have raised interest in its potential to similarly progress biological psychiatry. This study investigates the current utility of AI models in predicting psychiatric phenotypes in youth - a critical window for psychiatric diagnosis - using neuroimaging data from a large developmental clinical cohort. We assessed the predictive performance of machine learning models on diverse psychiatric and non-psychiatric targets. We show that while models are able to predict various targets from EEG and fMRI data, simple models using only readily-available factors such as demographics and recording site match their performance for clinical phenotypes. This pattern holds across clinical targets and replicates for state-of-the-art deep neural networks, suggesting that either neuroimaging data contains limited disease-specific information or that current methods cannot reliably identify such patterns. These benchmarking results provide important context regarding promises of modern AI in the field of biological psychiatry with a focus on youth.

## Introduction

Artificial intelligence (AI) holds immense promise to advance clinical workflows as a tool to analyze complex data such as brain dynamics, which are difficult for humans to interpret (Topol 2019). Psychiatry, with its diagnostic challenges due to heterogeneous symptoms and overlapping disorders, is a prime target for such innovations (Bzdok and Meyer-Lindenberg 2018; Schnack 2019). Here, AI can be used to discover objective biomarkers from brain recordings collected via functional MRI or electroencephalography (EEG). These non-invasive modalities are widely available and capture neural signatures linked to psychiatric conditions (Greicius 2008; Buzsáki and Watson 2012; Shor et al. 2023; Wen et al. 2025).

Yet, despite enthusiasm, the potential of applying AI to brain dynamics in psychiatry remains unclear (Varoquaux and Cheplygina 2022). While many studies report high diagnostic accuracy using neuroimaging data, these are often limited by small sample sizes, overfitting, and methodological flaws. Consequently, these models frequently fail to generalize to more realistic clinical settings (Chekroud et al. 2024; Marek and Laumann 2024). Comorbidity constitutes an important example of a mismatch with clinical settings: whereas studies commonly classify patients with a single diagnosis and healthy controls, comorbidity is common in the real world. For psychiatric conditions, rates are estimated to exceed 25% and even 50% for internalizing disorders (Hasin et al. 2018; Kessler et al. 2005; Teesson et al. 2009). This highlights the need to account for the complexity of clinical populations by advancing and evaluating models in realistic, comorbid populations.

Another challenge and opportunity in applying AI to brain dynamics in psychiatry lies in the developmental context. A majority of psychiatric disorders emerge before adulthood, making this a critical period for modeling (Kessler et al. 2005; McGrath et al. 2023). While this developmental window presents unique opportunities such as high neuroplasticity enabling more effective early interventions (Weyandt et al. 2020), it also poses diagnostic challenges to be solved. Symptoms can be fluid in youth (O’Connor et al. 2020) and psychiatric diagnosis is less reliable in children, which may result from increased difficulties in the articulation of their problems (Edelbrock et al. 1985; Hus and Segal 2021; Hancock et al. 2023). These factors together emphasize the need for objective, biological biomarkers for this population in particular.

A particularly striking observation is that studies with greater sample sizes tend to report worse rather than better predictive performance (Ioannidis 2005; Button et al. 2013; Arbabshirani et al. 2017; Flint et al. 2021; Kambeitz et al. 2018; Schnack and Kahn 2016; Winter et al. 2024), suggesting that the numerous smaller studies overestimate predictability. As noted in the cited works, methodological issues causing data leakage and low reliability of results contribute to this issue. Yet, academic pressure for novel and positive results further exacerbates this, incentivizing for iterative model and pipeline optimization on small datasets. This may result in overfitting to dataset-specific patterns rather than learning general biological patterns, which inflates prediction scores while hurting performance on external data. Publications introducing novel methods are particularly incentivized to yield ‘state of the art’ performance, rather than solely providing an unbiased evaluation.

The aforementioned issues contribute to the critical question of whether neuroimaging data provide substantial predictive value beyond simple and readily available variables such as age and sex, which are known to influence psychiatric outcomes (Alegría et al. 2018; McGrath et al. 2023; Sialino et al. 2021; Yang et al. 2024). Moreover, when neuroimaging models are found to be predictive, it is possible they rely on demographic or other factors such as recording site or head motion (Power et al. 2012; Rane et al. 2022; Snoek et al. 2019). Research relying on maximizing predictive performance often neglect these potential confounding effects, focusing solely on performance. However, this may lead to the false inference that psychiatric phenotypes can be predicted based on brain dynamics, rather than such confounding factors. Altogether, unbiased benchmarking with consideration for potential confounds is crucial to obtain an accurate understanding of the status quo of neuroimaging-based phenotype prediction in psychiatry.

Here we address these challenges and provide systematic analyses using the Healthy Brain Network (HBN) cohort (Alexander et al. 2017), a large (4000-participant) developmental cohort. It captures transdiagnostic psychiatric disorders with significant and realistic comorbidity rates reflective of the fluid symptoms and diagnostic complexities of early-onset mental health conditions. We systematically investigate the benefit of neuroimaging data by objectively benchmarking the predictive performance of AI models across a range of targets.

## Methods

### Participants

To enable predictive modeling on a broad range of psychiatric targets with high sample sizes, this study utilized multimodal neuroimaging data from the large-scale Healthy Brain Network (HBN) project (Alexander et al. 2017). The HBN targets a developmental population of children and young adults (5-21 years) and includes high rates of psychiatric comorbidity (65% with 2 or more diagnoses), reflective of clinical populations (Table 1). We provide further descriptive statistics in Figures A5, A6.

**Table 1.** Overview of the categorical prediction labels used for classification (left) and the dimensional prediction labels used for regression (right)

| Categorical Prediction Labels |  |  |  | Dimensional Prediction Labels |  |  |  |
| --- | --- | --- | --- | --- | --- | --- | --- |
| Label | n<sub>EEG</sub> | n<sub>fMRI</sub> | Description | Label | n<sub>EEG</sub> | n<sub>fMRI</sub> | Description |
| Clinical - Diagnosis |  |  |  | Clinical - Symptom severity |  |  |  |
| CGAS | 850 | 374 | Children's Global Assessment Scale total score; a cross-diagnostic measure of general psychological and social functioning. (Scored 1-100, classes: score <51 and >72) | CBCL | 2530 | 924 | Child Behavior Checklist, cross-diagnostic measure of behavioral/emotional problems in children, total score |
| ADHD | 2682 | 1132 | Attention deficit hyperactivity disorder diagnosis | SWAN | 2600 | 930 | Attention and behaviour regulation in ADHD, total score |
| ASD | 2682 | 1132 | Autism spectrum disorder diagnosis | ASSQ | 2647 | 1106 | Autism Spectrum Screening Questionnaire total score |
| Anxiety | 2682 | 1132 | Anxiety disorders diagnosis | SRS | 2531 | 919 | Social Responsiveness Scale total score: a measure of social impairment in autism |
| COM | 2682 | 1132 | Communication disorders diagnosis | SCARED | 1867 | 797 | Screen for child anxiety related disorders total score |
| Depression | 2682 | 11328 | Depression diagnosis | CELF | 2625 | 927 | Clinical Evaluation of Language Fundamentals total score |
| SLD | 2682 | 1132 | Specific learning disorders diagnosis | MFQ | 1814 | 990 | Mood and Feelings Questionnaire total score |
| Disruptive | 2682 | 1132 | Disruptive behavior disorders diagnosis | ARI | 2610 | 929 | Affective Reactivity Index total score: a measure of irritability and mood dysregulation |
| No | 2682 | 1132 | No diagnosis | WISC Full Scale IQ | 2219 | 814 | Full-Scale Intelligence Quotient via Wechsler Intelligence Scale for Children |
| Other | 2682 | 1132 | Diagnoses that do not fit others | Demographic |  |  |  |
| Demographic |  |  |  | Age | 2682 | 1219 | Age of participant at the time of scanning |
|  |  |  |  | Physical fitness and lifestyle |  |  |  |
|  |  |  |  | SDS | 2298 | 1053 | Sleep disturbance scale total score. |
| Sex | 2736 | 1219 | Biological sex of the participant | Upper Body Strength | 2229 | 818 | Average performance on upper body exercises from the Fitness Gram questionnaire |
| Income | 1677 | 763 | Parental income operationalized as an extreme group prediction task (low-income vs high income) | BMI | 1978 | 793 | Body mass index |
|  |  |  |  | Physical activity | 1489 | 753 | Total frequency of participation in vigorous physical activities in the past week |

### Neuroimaging Data

We use the EEG data of up to n=2682 and fMRI data of up to n=1219 (after exclusion based on head motion using mean frame-wise displacement (FD) > 0.5mm or max FD > 5mm). Standard preprocessing is used for EEG relying on the MNE Python package (Gramfort et al. 2013), which yielded 10-second epochs. fMRI data is preprocessed using the standard fmriprep pipeline (Esteban et al. 2019), but with normalization to age-specific MNI pediatric templates (age ranges 4–7, 7–10, 10–13.5, 13.5–21 years). Subsequently, we extract region-of-interest-based (ROI) timeseries from the 4D BOLD signal using the Brainnetome atlas consisting of 210 cortical and 36 subcortical parcels (Fan et al. 2016) using Nilearn (Abraham et al. 2014). Full details on our data preprocessing and participant exclusion criterion can be found in the appendix.

### Prediction Targets

We leverage the HBN’s rich phenotyping effort to construct a broad set of prediction targets, spanning cognitive, physical, and demographic dimensions in addition to both psychiatric diagnostics and clinical severity scores. First, we predict psychiatric diagnoses which were determined through clinical consensus using the Schedule for Affective Disorders and Schizophrenia—Children’s version (KSADS) (Alexander et al. 2017; Kaufman et al. 1997). We grouped diagnoses into DSM-based groups as suggested by (Langer et al. 2022) (Table 1). We additionally included sex, household income, and an aggregate measure of impairment (Children’s Global Assessment Scale; CGAS (Bird et al. 1987)); that we operationalized as binary groups representing extreme categories (Gijsen and Ritter 2025b). This resulted in a total of 12 binary prediction targets.

Given concerns about shortcomings of using discrete disease categories, and to align with the emerging view of psychopathology as a multi-dimensional continuum (Haslam et al. 2020; Hengartner and Lehmann 2017), we also predicted a comprehensive set of fourteen continuous variables, representing various clinical, cognitive, lifestyle and demographic dimensions (Table 1).

### Predictive Design

To investigate the predictability of psychiatric phenotypes, we break our analyses into two components. First, we establish a baseline of predictability by fitting machine learning models to neuroimaging data and directly predicting binary and continuous labels without controlling for confounds. While these analyses estimate predictability per se, they do not tell us whether disease-specific brain activity has predictive utility. Indeed, models may estimate factors such as sex, age, head-motion, or recording site from neuroimaging data and rely on them for clinical prediction. To investigate whether brain data improves prediction beyond such simple-to-acquire confounds, we perform a contrastive analysis. Specifically, the performance of models trained only on the aforementioned confounds is compared with models receiving both confounds and neuroimaging data as input features. We use subject sex, age, the modality-specific recording site, the data release number a participant first appeared in, as well as head motion (fMRI only) as the confound set.

### Machine Learning Models

To understand whether modern deep neural networks (DNNs, deep learning; DL) perform differently from traditional machine learning (ML) models in terms of predictive performance, we include and compare three approaches from each modeling paradigm. For the ML approach, we rely on a widely used and successful machine learning pipeline. This relies on feature extraction which constitutes a pre-specified data representation based on expert knowledge. These features are used to train a linear or more expressive, non-linear machine learning algorithm: linear support vector machine (SVM), non-linear SVM (radial basis function kernel), and a tree-based approach (XG-BOOST; (Friedman 2001; Chen 2016)). For ML models, we use a nested cross-validation approach with three ‘inner’ folds used for determining the regularization strength as well as performing feature selection (selecting 1%, 3%, 10%, 33%, or 100% of all features). The aforementioned five ‘outer’ folds are subsequently used to estimate test accuracy on unseen data without data leakage.

For fMRI feature extraction, models use functional connectivity between ROIs by computing their pairwise Pearson correlations. Meanwhile, for EEG we extract a broad set of features based on time-frequency analysis and information theory (please see the Appendix and (Engemann et al. 2022) for complete details). Both are in accord with established practices in the literature (Saeidi et al. 2021; Liu et al. 2025).

In contrast to the ML models, DL models can operate directly on the neuroimaging timeseries and learn to extract non-linear features during the training process and are thereby more flexible. We choose DNNs proposed specifically for the neuroimaging domain and which have shown success in previous work. For both modalities we include a convolutional neural network architecture (CNN) and a transformer-based model for timeseries (Bedel et al. 2023; Wang et al. 2024; Gijsen and Ritter 2025b, 2025a). For EEG, we additionally include a popular hybrid model combining a CNN and recurrent neural network (Li et al. 2022). For fMRI, we also include a popular graph-transformer operating on the functional connectivity values (Kan et al. 2022). Given computational constraints, we used conventional five-fold cross-validation for DL models. We therefore adopt the hyperparameters proposed by the original authors, which we note are in turn based on empirical neuroimaging analyses.

For all analyses we use a five-fold cross-validation scheme while stratifying on age, sex and the prediction target to obtain variability in model performance. To maximize sample sizes and maintain realistic comorbidity levels, we perform one-versus-rest prediction for the diagnostic labels. Additionally, we perform two control analyses. First, to investigate the effect of label imbalance, we repeat our analyses in a one-vs-one setting (diagnosis-vs-no diagnosis) using a smaller, balanced sample obtained by stratified undersampling, where each diagnostic group is approximately matched to the group without diagnoses based on demographics. Second, we repeat our analyses using a stricter head-motion inclusion criterion for fMRI data, lowering the threshold for mean FD from 0.5 to 0.2mm and max FD from 5mm to 1mm. Details on both aforementioned experiments can be found in the appendix.

## Results

We first evaluate models trained using neuroimaging data only (Figure 1A). We report on the area under the receiver operating characteristic curve (AUC; with 0.5 indicating chance-level performance) and Pearson correlation coefficients (*r*) for the best performing model. The highest scores are seen for demographic factors, namely age (*r*_*EEG*_ = 0. 855 ± 0. 009, *r*_*FMRI*_ = 0. 752 ± 0. 017) and sex (*AUC*_*EEG*_ = 0. 944 ± 0. 003, *AUC*_*FMRI*_ = 0. 699 ± 0. 038). Further non-psychiatric variables were predictable but to a lesser extent, including physical variables BMI (*r*_*EEG*_ = 0. 586 ± 0. 033, *r*_*FMRI*_ = 0. 351 ± 0. 064) and upper body strength (*r*_*EEG*_ = 0. 420 ± 0. 027, *r*_*FMRI*_ = 0. 260 ± 0. 053), and parental income (*AUC*_*EEG*_ = 0. 653 ± 0. 016, *AUC*_*FMRI*_ = 0. 552 ± 0. 018). Psychiatric variables which had highest predictive scores were language proficiency (CELF scale; *r*_*EEG*_ = 0. 488 ± 0. 020, *r*_*FMRI*_ = 0. 390 ± 0. 034) and depression diagnosis (*AUC*_*EEG*_ = 0. 731 ± 0. 017, *AUC*_*FMRI*_ = 0. 704 ± 0. 044). While some diagnostic or continuous prediction scores were very close to *AUC* = 0. 5 or *r* = 0. 0, most clinical targets appeared mildly predictable with *AUC*≅0. 6 or *r*≅0. 1 for at least one neuroimaging modality.

**Figure 1.**
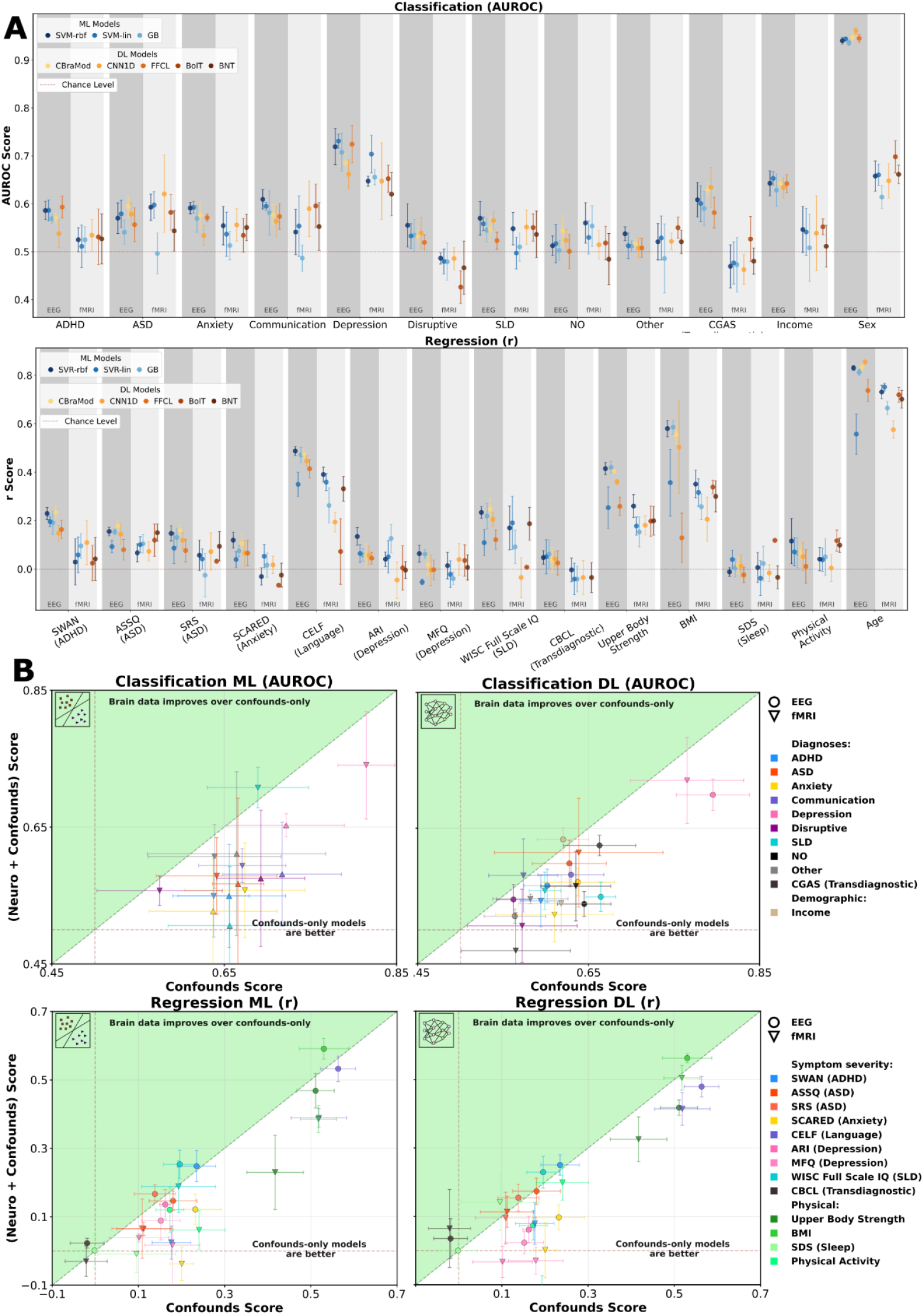
**(A)** Predictive performance using neuroimaging data with traditional machine learning (ML) and modern deep learning (DL) models for binary classification targets (top) and continuous targets (bottom). The horizontal dotted line indicates 0.5 AUROC (chance-level performance; top) and r=0 (no linear correlation between targets and predictions; bottom). **(B)** Performance comparison between models using only simple confound variables as input and models using both confounds and neuroimaging data. The diagonal indicates relative performance, with confound-only models performing worse (above diagonal; green), equal (on diagonal), or better (below diagonal). Plots are separated between ML and DL models (left-right) and classification and regression targets (top-bottom). Error bars indicate the standard deviations across five different subject splits.

Crucially, to investigate whether machine learning models learned disease-specific brain patterns, we contrast prediction scores for models relying only on a small number of confound variables compared to operating on both confound variables and brain data (Figure 1B). We observe that for almost none of the targets model predictions are clearly improved when additionally learning from neuroimaging data. While mean predictability for some clinical targets is slightly higher when neuroimaging data is included, variability over subject-splits prevents drawing conclusions of marked improvements. Some non-psychiatric predictions were improved to a greater extent by including EEG data, although still with notable variability across subject-splits: IQ-prediction (WISC; *r*_*EEG*+*CON*_ = 0. 253 ± 0. 042, *r*_*CON*_ = 0. 196 ± 0. 038), BMI-prediction (*r*_*EEG*+*CON*_ = 0. 591 ± 0. 030, *r*_*CON*_ = 0. 530 ± 0. 057), and parental income prediction (*AUC*_*EEG*+*CON*_ = 0. 681 ± 0. 025, *AUC*_*CON*_ = 0. 626 ± 0. 042). Similar patterns are observed when using stricter head-motion inclusion criteria (Appendix Figure A1).

Contrasting traditional machine learning pipelines and modern deep neural networks, we do not observe major differences in predictive performance despite operating on significantly different input data and a variety of tested model architectures. The best performing model family is inconsistent across prediction targets, with generally small score discrepancies (less than 0.04 AUC or *r*) for both binary and continuous contexts.

We we conduct one-sided paired t-tests comparing neuro+confounds versus confounds-only performance across clinical targets, separately for each combination of modality (EEG, fMRI), model family (ML, DL), and target type (classification, regression), yielding 8 tests. None reached significance (all corrected *p*_*FDR*−8_ > 0.05), indicating that adding neuroimaging features did not significantly improve prediction over confounds alone. Finally, we compute effect sizes for the associations between confound variables and prediction targets. We observe significant relationships across many variables and present these results in Appendix Figure A4.

## Discussion

This study investigated whether AI models can leverage neuroimaging data from EEG and fMRI in youth, a developmental window of particular clinical relevance. To this end, we leveraged a clinically heterogeneous developmental cohort to improve the prediction of psychiatric phenotypes beyond simple factors such as demographics, dataset meta-data, or head-motion. Our primary finding is that while machine learning models accurately predicted demographic factors such as age and sex from brain data, they did not significantly improve predictive performance for psychiatric targets - whether categorical diagnoses or continuous measures - compared to confound-only models. This pattern persisted across data modalities, a range of model architectures (from linear models to bespoke deep neural networks), and a large set of clinical targets.

These results point to two potential explanations: either resting-state neuroimaging data contains limited disease-specific information, or current machine learning models are unable to extract such information. Given the importance of sample size for machine learning, it is important to note that even larger datasets might be necessary for models to reliably extract such forms of predictive information. The observation that models successfully learned between-subject variability predictive of demographics suggests they are capable of fitting generalizable patterns in the data. Moreover, repeating fMRI analyses with stricter head-motion inclusion criteria (mean FD < 0.2mm, max FD <1mm) did not considerably alter the results, indicating that poor data quality is unlikely to be the primary driver. Therefore, our analyses indicate less disease-specific signals in the neuroimaging data than expected, which appears to apply to both categorical and dimensional targets.

Indeed, previous work suggests that models trained on neuroimaging data do not see performance saturation even on datasets significantly larger than considered here (Kiessner et al. 2023; Schulz et al. 2020). This is especially problematic for the psychiatric domain: whereas large datasets are being collected for fMRI (Bamberg et al. 2024; Miller et al. 2016) and EEG (Obeid and Picone 2016), they lack extensive clinical phenotyping or diagnosed psychiatric patients. The problematic data-hungry property of modern machine learning is unlikely to be overcome with greater sample sizes. Instead, it may rather indicate a need for methodological innovations to reliably extract relevant information, to the extent that it is present. Yet, it may furthermore be necessary to overcome issues pertaining diagnostic label noise as well, which further complicates psychiatric applications. These challenges may be especially pronounced in youth, due to factors such as the fluidity of symptoms and difficulties with expressing symptoms, potentially amplifying label noise.

Our findings are at once in contrast to numerous smaller studies reporting high diagnostic accuracy using neuroimaging while simultaneously corroborating larger studies and reviews observing diminished predictive performance as sample sizes increase (Arbabshirani et al. 2017; Flint et al. 2021; Kambeitz et al. 2018; Schnack and Kahn 2016). Such discrepancies suggest that methodological issues in the literature as well as scientific publication incentives may have led to inflated predictability estimates in prior work. By using a large, comorbid sample with limited prior machine learning analyses, our study attempted to reduce such biases, offering a more realistic benchmark for AI-based psychiatric prediction. Our results also highlight the importance of accounting for the role of confounds if researchers aim to infer on disease-specific brain dynamics. This is highlighted by the numerous significant associations between confounds and prediction targets of interest. Indeed, benchmarking against confounding variable models is crucial for understanding the additive value of neuroimaging models generally, as exemplified by our depression classification results. To this end, the contrastive analysis approach employed here may also be complemented with or substituted for other methods such as propensity score matching or counterbalancing (Rosenbaum and Rubin 1983; Rane et al. 2022; Schnack and Kahn 2016; Snoek et al. 2019).

Some limitations of the current work deserve mention. The investigated pediatric sample may be of lower data quality in ways other than head-motion, which could impair ML performance. As we trained a range of prediction models spanning classical ML to modern deep neural networks, computational constraints precluded extensive hyperparameter tuning. Nevertheless, the risk of poor model fit was mitigated by confirming model convergence during training. Finally, our analysis only concerns supervised approaches. Future work may include methodologies currently being developed which allow for the pretraining of models on large, unlabeled datasets, before being applied to smaller, clinical datasets. This may help overcome the data-hungry nature of current methods.

## ACKNOWLEDGEMENTS

This research was funded by Gemeinnützigen Hertie-Stiftung and the Deutsche Forschungsgemeinschaft (DFG) through FOR 5187 (project number 442075332). Additional support was provided by the Machine Excellence Cluster and DFG through the Germany’s Excellence Strategy (EXC 2064 - project number 390727645) and the following projects: CRC 1404 (project number 414984028), TRR 265 (project number 402170461), and RU 5363 (project number 459422098).

None of the authors have any financial disclosures or conflicts of interests.

## APPENDIX

### fMRI Preprocessing and inclusion criterion

Subjects with functional resting-state, anatomical and field map data were selected for preprocessing (n=1920), an exclusion criterion of 0.5 mm mean FD and 5 mm max FD was used for our main analysis (n=1219), a more strict exclusion criterion of 0.2 mm mean FD and 1 mm max FD was used for our motion control analysis (n=497). Preprocessing was performed using fMRIprep (Esteban et al., 2019), which included skull stripping, brain extraction, tissue segmentation, and surface reconstruction for anatomical data, and non-steady-state frames removal, head motion correction, slice time correction, susceptibility distortion correction (SDC) for functional data. We used age-specific MNI pediatric templates for the spatial normalization step to account for the variability of brain sizes in this developmental cohort (age ranges: 4-7, 7-10, 10-13.5, 13.5-21).

For postprocessing using Nilearn, we applied a frame-wise displacement threshold of 0.5mm to censor frames with high motion (a process known as scrubbing) as suggested by (Power et al. 2012). This process ameliorates the systematic local spurious correlations caused by head motion. The first 365 uncensored volumes were used for subjects with more than 365 volumes. Deep learning models require continuous fMRI data, so for these models the scrubbing step was omitted while excluding four additional subjects as they had less than 365 total volumes. Further preprocessing steps included: 24 parameter motion regression (Friston et al. 1996), ROI timeseries extraction using the Brainnetome atlas (210 cortical and 36 subcortical parcellations) (Fan et al. 2016), and Butterworth bandpass filtering using a 0.009 to 0.08Hz cutoff.

#### EEG Feature Extraction

Following the methodology of (Engemann et al. 2022), we extracted a feature set from the EEG data: this set spanned time, frequency and time-frequency domain features, statistical features, as well as power and information-theory based features. This resulted in 6448 features per subject. Specifically, the following features were computed for individual channels, concatenated across channels, and finally we selected the median for each feature across epochs: standard-deviation, kurtosis, skewness, quantiles (10%, 25%, 75%, 90%), peak-to-peak amplitude, mean, power ratios among frequency bands (0-2, 2-4, 4-8, 8-13, 13-18, 18-24, 24-30, 30-49Hz), spectral entropy, approximate and sample entropy, temporal complexity, Hurst component, Hjort complexity and mobility, line length, energy of wavelet decomposition coefficients, Higuchi fractal dimension, number of zero crossings and the per-channel SVD Fisher information.

#### Details on Machine Learning Pipeline

For ML methods, we follow a well-established pipeline. Specifically, for feature normalisation we used z-scoring (with parameters estimated only on the training data to prevent data leakage). For feature selection using scikit-learn, we first use a variance threshold to remove any zero-variance features, followed by the SelectFromModel method. The latter involves training a proxy model on the data to estimate the importance of each feature. This allows for the selection of a subset of the most informative features before training the final classifier. We account for class imbalance by down-weighting the majority class in the loss function.

Specifically, for the linear SVM, a LinearSVC estimator was used as the proxy, with feature importance determined by the magnitude of the learned coefficients. For the non-linear RBF SVM, a RandomForestClassifier served as the proxy model to derive feature importances. Finally, the XGBoost model used an XGBClassifier as its own proxy for feature selection. The number of features to select was treated as a hyperparameter, choosing from the top 1%, 3%, 10%, 33%, or 100% of features during the cross-validation process.

#### Deep Learning Models

We provide further details on the different deep learning models which were evaluated. Unless noted otherwise, we set hyperparameters to their recommended values by the original authors. For models using both neuroimaging data and confounds, we concatenate the confounds with the neuroimaging-extracted features prior to the task-specific prediction layers.

#### CNN (fMRI)

A 1D Convolutional Neural Network (CNN) The architecture consists of a hierarchical feature extractor with three sequential 1D convolutional layers. These layers progressively reduce the temporal dimension using a stride of 2 while increasing the feature map depth (from 64 to 128, then 256 channels) and using decreasing kernel sizes (7, 5, and 3, respectively). Following the final convolutional layer, a global average pooling operation is applied across the temporal dimension to produce a fixed-size feature vector representing the entire time series. This feature vector can optionally be concatenated with subject-level confounding variables. Finally, a fully connected (linear) layer maps the resulting vector to the output classes.

#### BrainNetworkTransformer (fMRI)

The Brain Network Transformer is an architecture specifically adapted for the unique properties of brain connectivity graphs. Instead of using complex positional encodings for the transformer, it uses the connection profile for each Region of Interest (ROI) as the initial node feature, a method that naturally encodes both structural and positional information suitable for the dense nature of brain networks. The model utilizes a standard multi-head self-attention mechanism, which learns predictive pairwise relationships without incorporating edge weights. Its primary architectural innovation is the Orthonormal Clustering Readout (OCRead) operator, a graph pooling function that generates a graph-level embedding by first performing self-supervised soft clustering of nodes. This process leverages an orthonormal projection to ensure the resulting cluster-aware embeddings are highly distinguishable, effectively capturing the brain’s inherent modular organization.

#### BolT (fMRI)

BolT is a transformer architecture designed to hierarchically analyze fMRI BOLD time series by segmenting the data into overlapping temporal windows. Its core innovation is the Fused Window Multi-head Self-Attention (FW-MSA) mechanism, which facilitates communication between adjacent temporal segments by allowing base tokens within a window to compute attention with fringe tokens from its neighbors. This operation is performed within a cascade of transformer blocks where the degree of window overlap is progressively increased, enabling the model to transition from learning local to global representations. For classification, BolT utilizes distinct classification tokens for each window and introduces a novel cross-window regularization loss to enforce consistency among these high-level feature representations across the entire time series. Finally, a token fusion step aggregates the multiple representations learned for each time point as it appears across different overlapping windows.

#### CNN (EEG)

A 1D residual Convolutional Neural Network (CNN) architecture, characterized by a multi-branch design to capture features across multiple temporal scales. The network begins with 32 learnable convolutions each for three different kernel lengths (4, 8, 16), followed by a series of four residual blocks. Each block’s main path employs a set of parallel 1D convolutional layers with different kernel sizes (also 4, 8, 16), whose feature maps are concatenated to learn simultaneous temporal patterns. A standard skip connection, using 1×1 convolutions to match dimensions, is added to the main path’s output before a final max-pooling layer performs temporal downsampling. After the last block, a global average pooling layer summarizes the features into a fixed-size vector, which can then be passed to a task-specific head, for which we use a 2-layer MLP with ReLU non-linearity.

#### FFCL (EEG)

A hybrid architecture designed for time-series classification that processes input data through two parallel branches. The first branch is a CNN that performs hierarchical feature extraction using two sequential convolutional blocks, where the second block is a depthwise separable convolution. Feature vectors are extracted after each of these two blocks to capture patterns at different levels of abstraction. The second branch consists of a Long Short-Term Memory (LSTM) network that processes the entire input sequence to model temporal dependencies, with its final hidden state serving as a sequential feature summary. In the final stage, the feature vectors from both levels of the CNN branch and the feature vector from the LSTM branch are concatenated. This fused representation is then passed through a fully connected layer to produce the classification output.

#### CBraMoD (EEG)

To adapt the transformer architecture for the unique structure of EEG data, CBraMod processes the signal by first creating single-channel patches and encoding them using parallel time-domain (convolutional) and frequency-domain (FFT-based) branches. The model’s core is a novel criss-cross transformer backbone that replaces the standard full attention mechanism with two parallel attention components designed to model spatial (inter-channel) and temporal (intra-channel) dependencies separately. To handle the variability in channel configurations across different datasets, CBraMod employs a convolutional network to dynamically generate positional embeddings, thereby enhancing the model’s adaptability to diverse EEG formats.

#### Effects of head-motion inclusion criteria on performance

The pediatric demographics of the HBN dataset make it a highly important cohort to study, while also introducing higher risk for excessive head motion. This, in turn, calls for consideration of the trade-off between sample size and data quality dictated by the head-motion inclusion threshold. To ensure that our findings are not driven by the threshold used in our main analyses, we repeated the analyses using a stricter threshold (mean FD < 0.2 mm, max FD < 1 mm). This yields reduced sample sizes more comparable to those used in the existing literature. Specifically, all diagnostic labels (ADHD, ASD, Anxiety, COM, Depression, Disruptive, SLD, NO, Other) had n=453; Age and Sex had n=497, CGAS n=146, Income n=298, ARI n=363, ASSQ n=440, BMI n=328, CELF n=359, CBCL n=353, MFQ n=418, SCARED n=320, SDS n=405, SRS n=351, SWAN n=360, WISC_FSIQ n=288, Phys_Activity n=304, UP_Strength n=310.

The results are shown in Figure A1. We observe similar predictive performance despite the stricter threshold. These findings therefore suggest that neither larger samples with lenient thresholds nor smaller samples with stricter thresholds enable clinical prediction beyond simple confound models.

**Figure A1.**
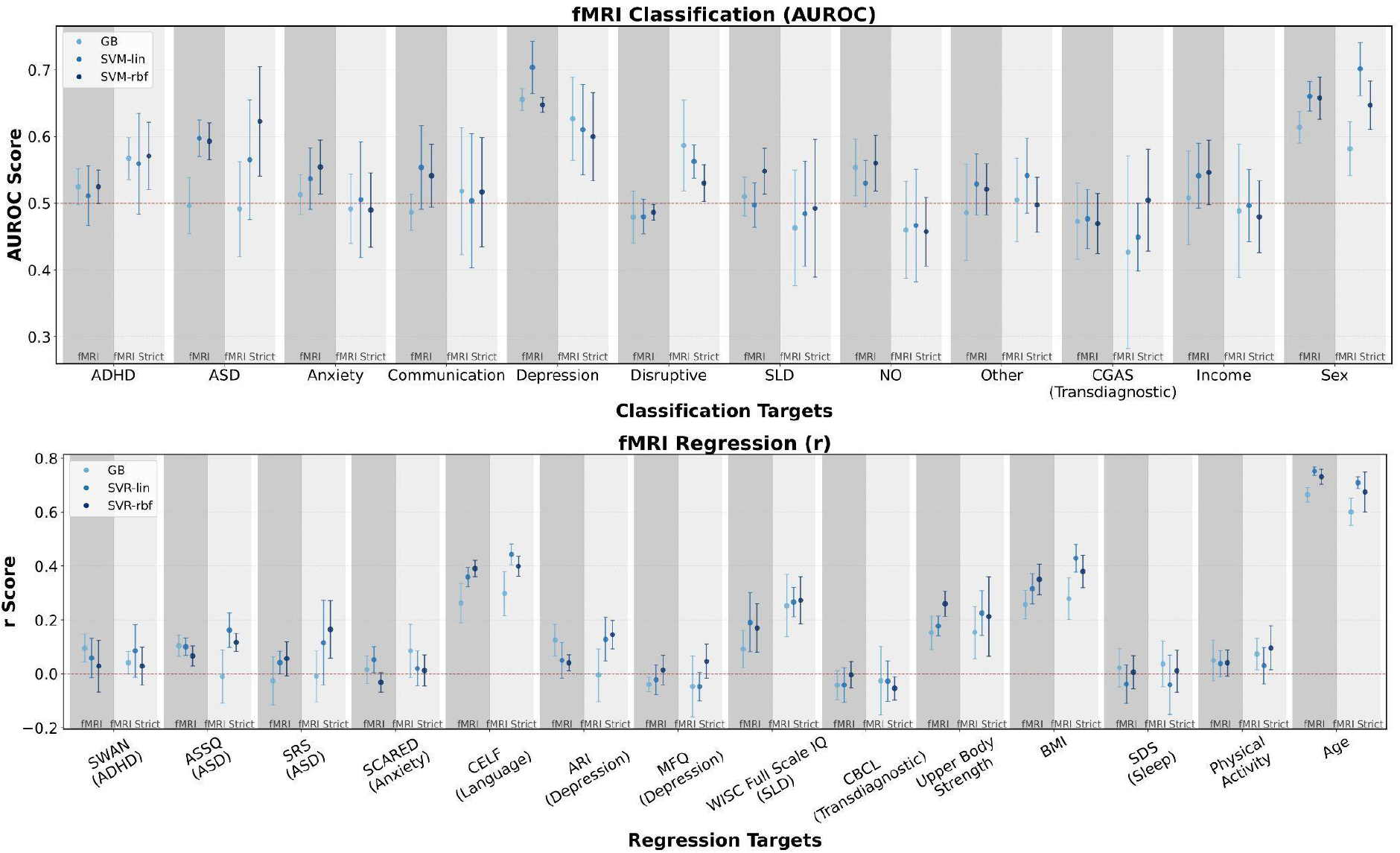
We evaluate predictive performance for binary targets (top) and continuous targets (bottom) using fMRI data following stricter inclusion criterion based on head motion. We observe no distinct differences in resulting performance capability.

#### One-vs-one classification

A one-vs-rest setup, in which a single diagnosis needs to be separated from all diagnostic classes, was prioritized for our main analyses as it enables larger sample sizes and is more realistic with greater relevance to clinical settings. However, because one-vs-one setup is more widely used in the scientific literature, we repeat our analyses using this approach. In a one-vs-one setup, a single diagnosis needs to be separated from the no-diagnosis group (e.g., subjects with only an ADHD diagnosis are contrasted with subjects without any diagnosis (“NO” diagnosis group)). Besides bridging the gap to a significant part of the literature, it provides insight as to whether our findings result from class imbalance. Specifically, we balance sample sizes between classes, rather than addressing it through optimization adjustments (i.e., down-weighting the majority class in the loss function). Finally, a one-vs-one setting may aid models in finding a better decision boundary between two classes due to reduced heterogeneity in the negative class.

We use stratified subsampling to obtain balanced sample sizes per class. Specifically, The no-diagnosis group serves as the negative class (N = 124 for fMRI, M = 270 for EEG). For each diagnostic group, we sample subjects with only that diagnosis as the positive class, matched as closely as possible to the no-diagnosis group in both demographics and sample size. This yields approximately equal numbers of positive and negative samples per diagnosis, with a few exceptions where exact matching of the strata was not possible due to limited availability of subjects belonging to a stratum: EEG depression (250), EEG COM (266), fMRI COM (123), and fMRI ASD (121).

The results are shown in Figure A2. We obtain a similar pattern as observed in our main analyses, suggesting that our findings are robust to variations in the formulation of the clinical prediction problem, rather than being specific to a one-vs-rest classification setup.

**Figure A2.**
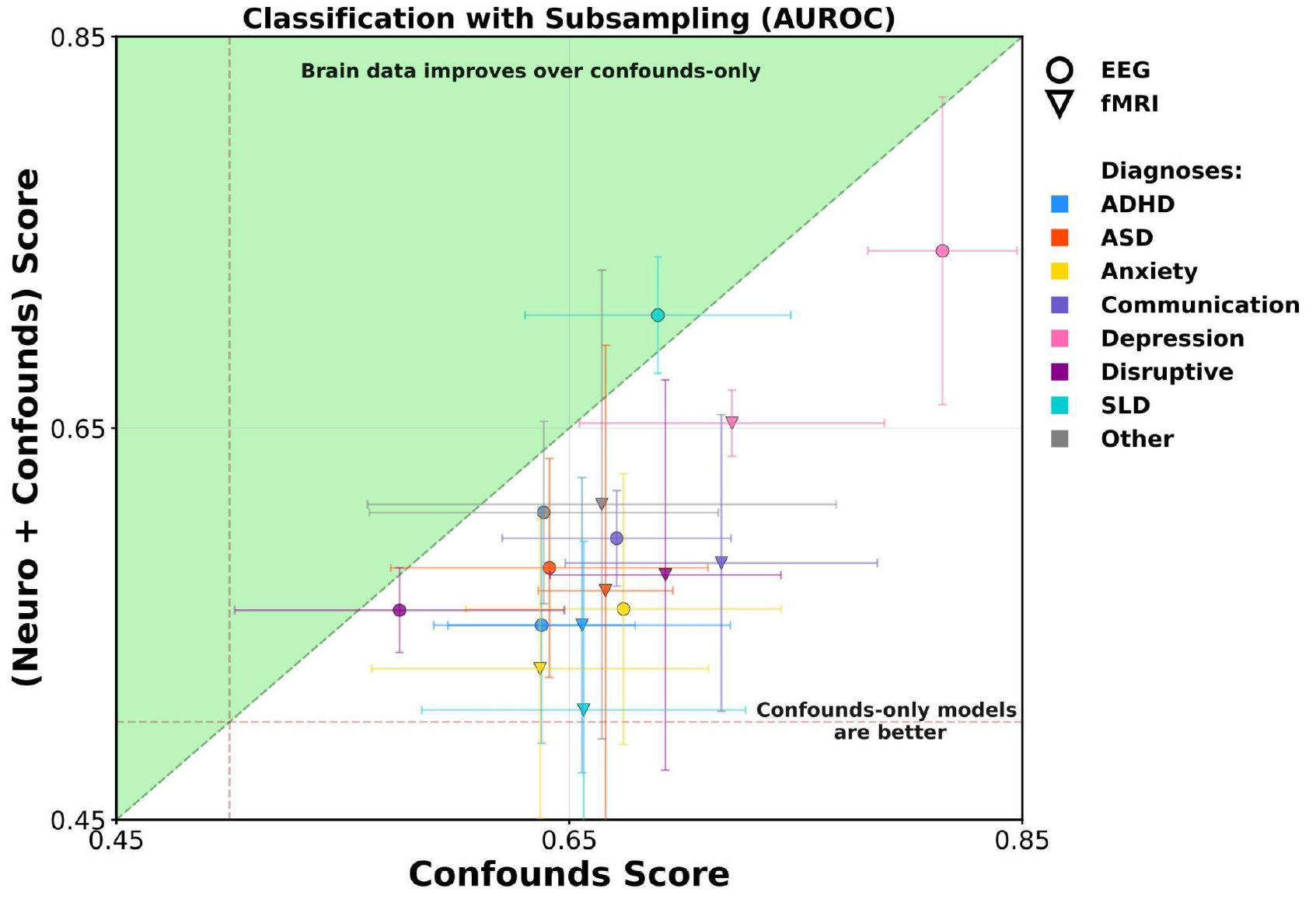
We repeat the comparative classification analyses using a one-vs-one classification setup, instead of performing one-vs-rest classification. We observe the same trend as in our main analysis as neuroimaging models do not reliably outperform confound-only models.

### Additional DL Models Results

**Figure A3.**
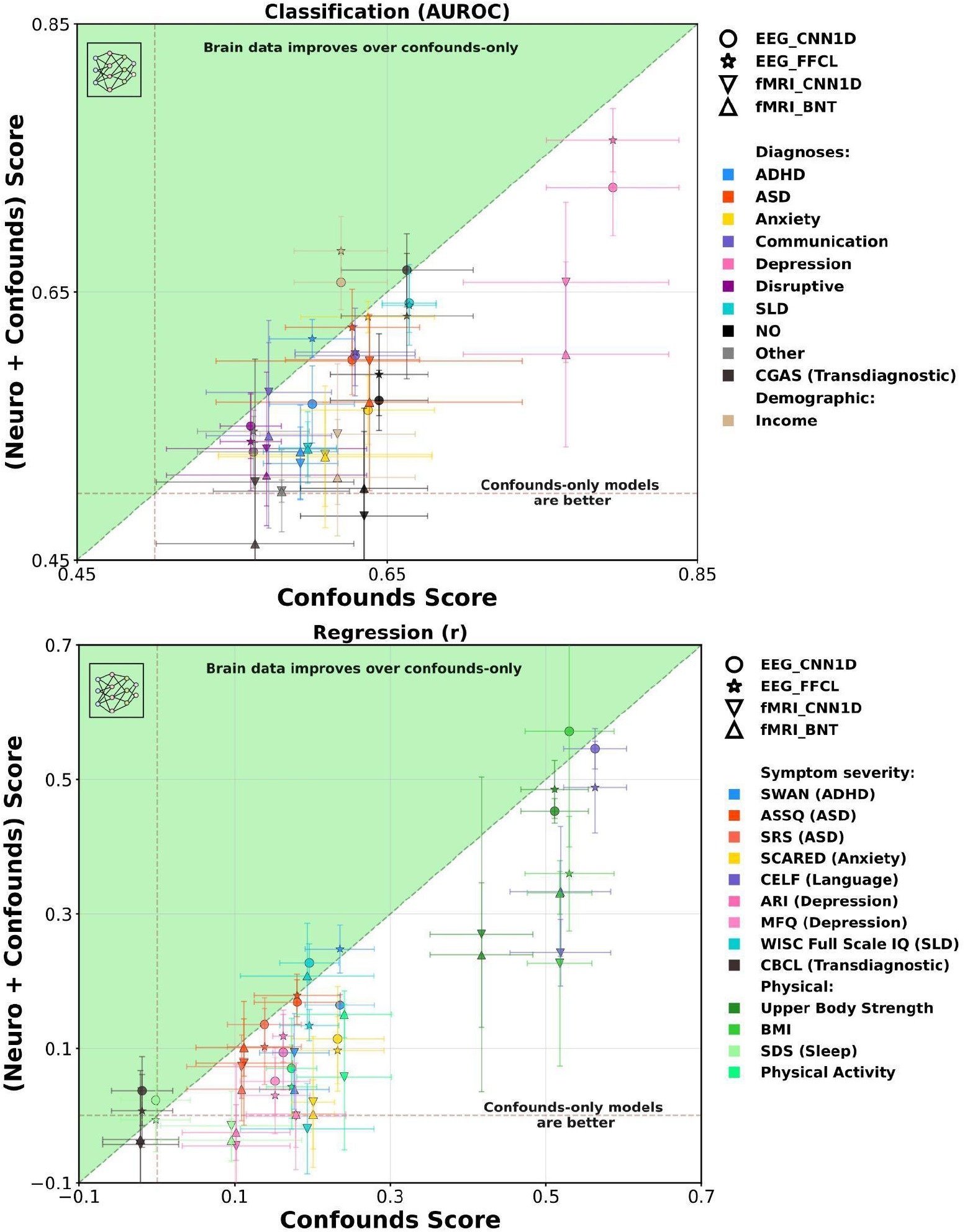
Whereas the main analysis presented the results of one deep learning (DL) model per modality for visual clarity, here we present the full results for the remaining DL models. Despite significantly different model architectures, no model enabled clearly improved performance using neuroimaging data in addition to simple-to-acquire confound factors.

### Confound Analyses

**Figure 4A.**
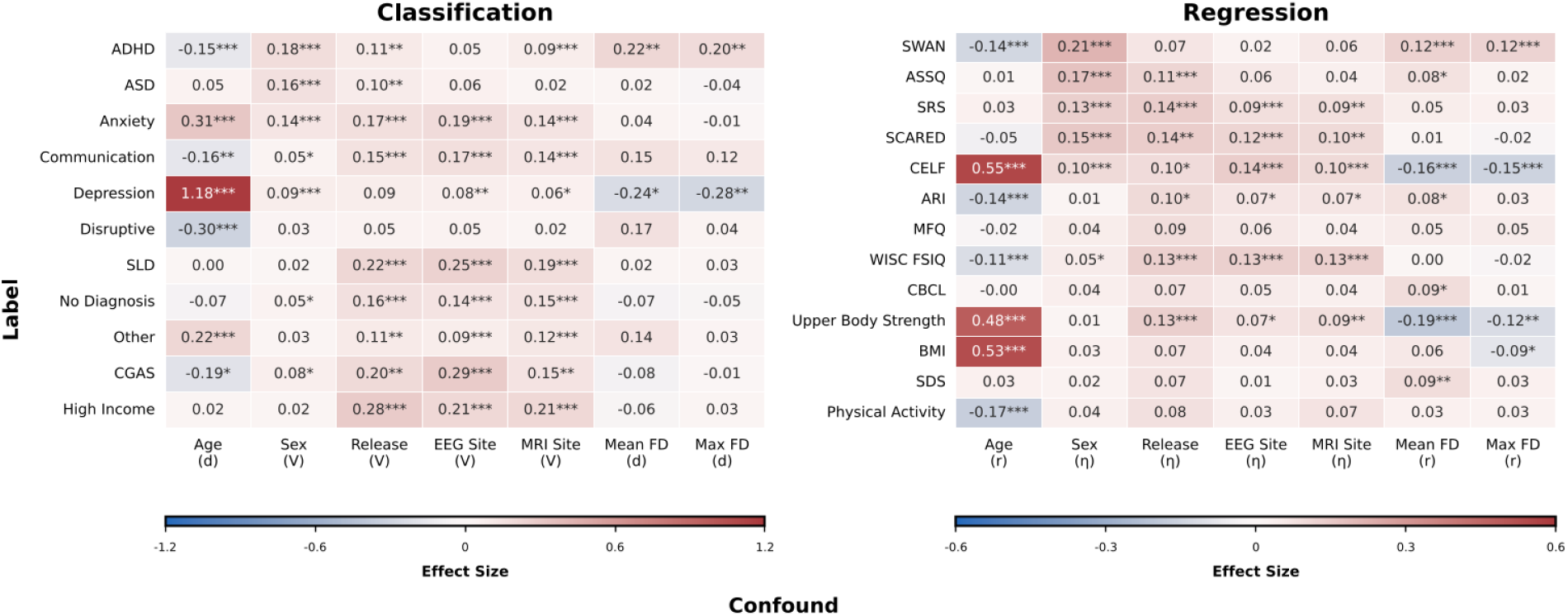
Effect sizes quantifying associations between prediction targets and potential confounds. Classification: Cohen’s d (continuous confounds) or Cramér’s V (categorical). Regression: Pearson r (continuous) or η^2^ (categorical). *p<0.05, **p<0.01, ***p<0.001, FDR-corrected (Benjamini-Hochberg). FD: framewise displacement, M: male, F: female

### Additional Dataset Statistics

**Figure 5A.**
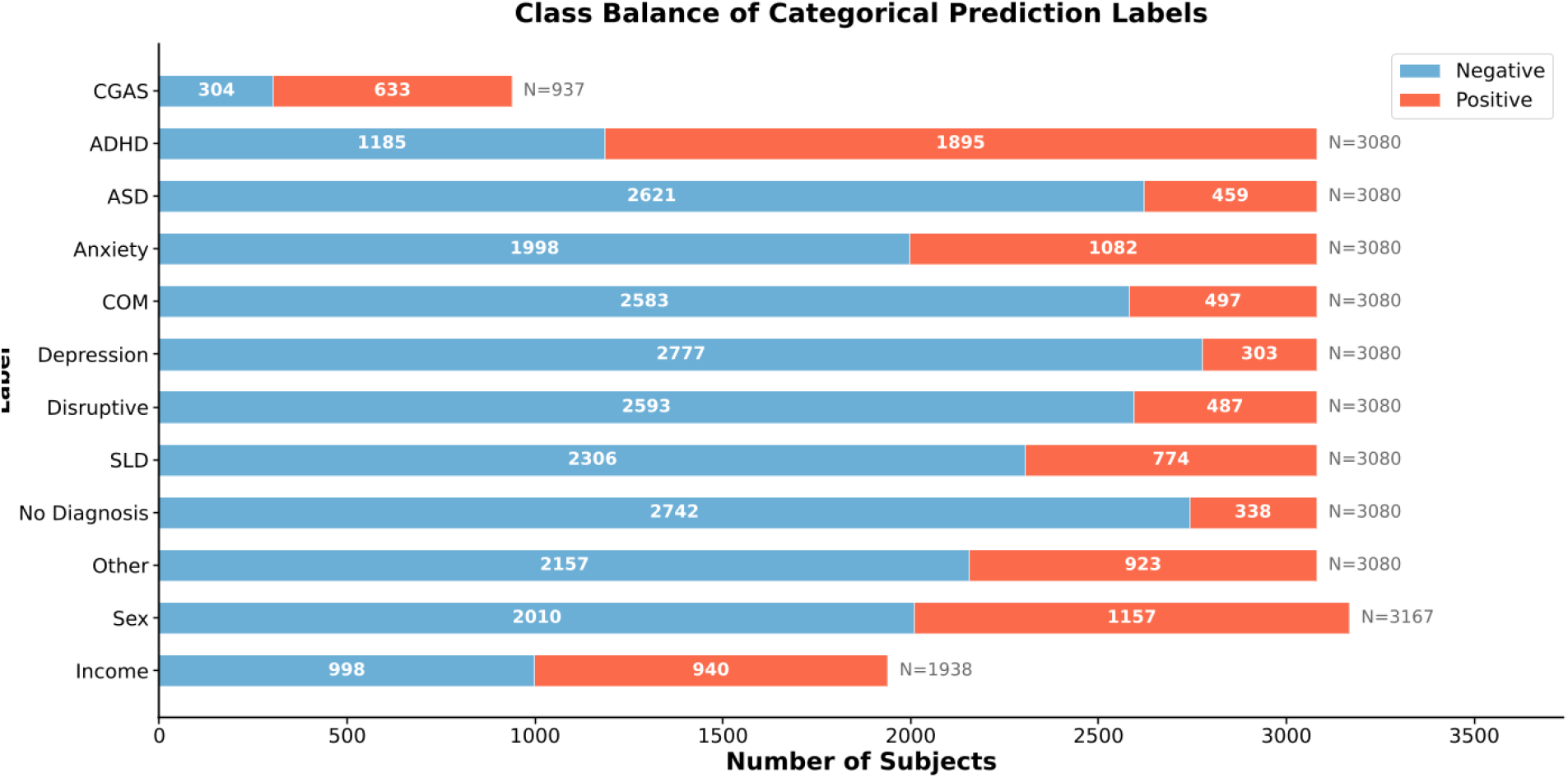
Class balance of categorical prediction labels across the combined EEG and fMRI dataset (union of both datasets, with duplicate subjects removed). Each bar shows the number of subjects in the positive and negative class for a given label. Labels are ordered as in Table 1. Total sample size per label (N) is indicated to the right of each bar.

**Figure 6A.**
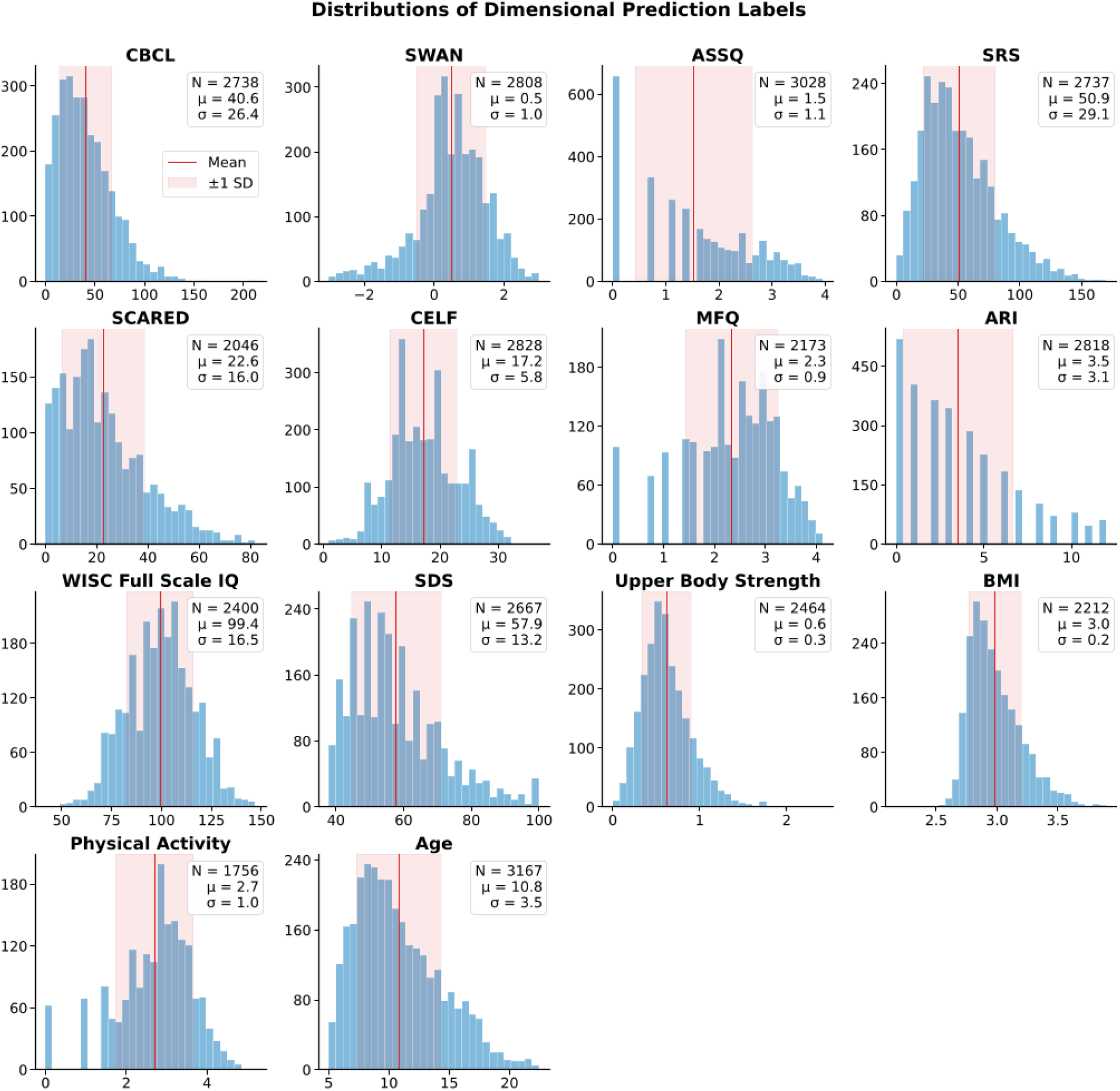
Distributions of dimensional prediction labels across the combined EEG and fMRI dataset (union of both datasets, with duplicate subjects removed). Each panel shows the histogram of scores (y-axis) for a given label on its original scale (x-axis). The red vertical line indicates the mean (µ), and the shaded region spans ±1 standard deviation (σ). Sample size (N), mean, and standard deviation are reported in each panel. Labels are ordered as in Table 1.

